# Acetylation of fission yeast tropomyosin does not promote differential association with cognate formins

**DOI:** 10.1101/2022.11.03.514925

**Authors:** Qing Tang, Luther W. Pollard, Kaitlin E. Homa, David R. Kovar, Kathleen M. Trybus

**Affiliations:** Department of Molecular Physiology and Biophysics, University of Vermont, Burlington VT; Molecular Genetics and Cell Biology, Biochemistry and Molecular Biology, the University of Chicago, Chicago, IL

**Keywords:** actin, tropomyosin, formin, myosin, acetylation, *S. pombe*

## Abstract

It was proposed from cellular studies that *S. pombe* tropomyosin Cdc8 (Tpm) segregates into two populations due to the presence or absence of an amino-terminal acetylation that specifies which formin-mediated F-actin networks it binds, but with no supporting biochemistry. To address this mechanism *in vitro*, we developed methods for *S. pombe* actin expression in Sf9 cells. We then employed 3-color TIRF microscopy using all recombinant *S. pombe* proteins to probe *in vitro* multicomponent mechanisms involving actin, acetylated and unacetylated Tpm, formins, and myosins. Acetyl-Tpm exhibits tight binding to actin in contrast to weaker binding by unacetylated Tpm. In disagreement with the differential recruitment model, Tpm showed no preferential binding to filaments assembled by the FH1-FH2-domains of two *S. pombe* formins, nor did Tpm binding have any bias towards the growing formin-bound actin filament barbed end. Although our *in vitro* findings do not support a direct formin-tropomyosin interaction, it is possible that formins bias differential tropomyosin isoform recruitment through undiscovered mechanisms. Importantly, despite a 12% sequence divergence between skeletal and *S. pombe* actin, *S. pombe* myosins Myo2 and Myo51 exhibited similar motile behavior with these two actins, validating key prior findings with these myosins that used skeletal actin.

## Introduction

A central open question in cell biology is how does the actin cytoskeleton self-segregate into functionally distinct F-actin networks subpopulations? Defining the chain of events that drive inter-network sorting from the initiation of assembly down to differential assortments of side-binding actin filament regulators is vital to understanding this question. The fission yeast *Schizosaccharomyces pombe* and other yeast systems have been studied extensively to help define core aspects of cytoskeletal self-organization, particularly because they possess a small number of distinct actin structures whose formation and function each involve a different collection of actin binding proteins (reviewed by (Kovar et al., 2011; Suarez and Kovar, 2016)). In yeast, endocytic patches are composed of branched actin networks, whereas the contractile ring and actin cables utilize linear actin structures. The mechanisms that occur downstream of actin nucleation and which affect the differential recruitment of actin binding proteins are just beginning to be elucidated. It has been recently shown that α-actinin and tropomyosin synergistically cooperate to compete with fimbrin and ADF/cofilin and vice versa for binding to filament sides to drive self-sorting networks *in vitro* (Christensen et al., 2017; Christensen et al., 2019). On the other hand, tropomyosin becomes essential for formin-based cable assembly when cofilin is enhanced by AIP1 (Okada et al., 2006; Pollard et al., 2020). Together, these observations suggest that network self-sorting occurs as a cumulative function of assembly-disassembly kinetics and cooperative reinforcement feedback by teams of actin side-binding proteins. Importantly, the possibility that the nucleators themselves could provide a direct positioning signal for downstream actin binding proteins has been raised.

Linear actin structures are nucleated and elongated by different formins in fission yeast. The contractile ring, which separates the daughter cells at the end of mitosis, is assembled by Cdc12, while the actin cables assembled by For3 drive polarization by serving as the track along the length of the cell for cargo delivery towards the two cell poles (Bendezú and Martin, 2011; Feierbach and Chang, 2001). It is unclear how actin-binding proteins differentially associate with these two distinct linear structures, both of which are stabilized by the sole and essential fission yeast tropomyosin Cdc8 (Balasubramanian et al., 1992; Kovar et al., 2011; Michelot and Drubin, 2011). One hypothesis suggests that different formins, through direct interaction with tropomyosin, control the composition of proteins bound to actin filaments downstream (Coulton et al., 2010; Johnson et al., 2014; Skoumpla et al., 2007). According to this proposed mechanism, tropomyosin Cdc8 is segregated into isoform-like populations by the presence or absence of an amino-terminal acetylation that specifies which formin it binds. If true, such a mechanism would have a profound impact on how actin networks self-segregate to perform different cellular functions, and may explain how tropomyosin isoforms (> 40 isoforms in mammalians) are sorted into many different actin networks in higher eukaryotes (Gunning et al., 2015). Additionally, higher eukaryotes have non-redundant actin isoforms (Perrin and Ervasti, 2010), which may influence tropomyosin sorting. To date, the tropomyosin sorting mechanism by formin has not been fully reconstituted *in vitro* with all native components.

Previous methods for fission yeast actin purification produced actin with high purity, but limited yield (Takaine and Mabuchi, 2007; Ti and Pollard, 2011). An alternative method for actin expression in a heterologous system, utilizing an actin-thymosin fusion construct, has recently allowed purification of mg quantities of recombinantly expressed actin (Funk et al., 2019; Lu et al., 2015; Lu et al., 2016; Noguchi et al., 2007). Here we applied a similar expression strategy to obtain pure fission yeast actin using the *Sf*9/baculovirus expression that yields tens of milligrams of *S. pombe* actin. By combining components from the same species *in vitro*, we found that the FH1-FH2 domains of fission yeast formin isoforms do not specify the tropomyosin variant (acetylated versus unacetylated) that associates with the actin filaments, and the sorting of tropomyosin was primarily determined by the apparent affinity between actin and tropomyosin. This observation suggests that additional molecular components are likely required for the tropomyosin sorting mechanisms proposed from cellular experiments (Johnson et al., 2014). We also found that the motor activities of two recently characterized fission yeast myosins (Myo2 and Myo51) are similar when either skeletal or fission yeast actin is used, unlike budding yeast actin which functions most effectively with its cognate myosins (Stark et al., 2011). Our study extends the ability to examine *in vitro* current hypotheses in cytoskeleton organization using a full complement of recombinant *S. pombe* proteins.

## Results

### Expression and purification of recombinant *S. pombe* actin in Sf9 cells

To express *S. pombe* actin using the baculovirus/insect cell expression system, we used a published strategy in which the actin monomer-sequestering protein human thymosin-β4-His_6_ is cloned onto the C-terminus of actin following a linker (Lu et al., 2015; Lu et al., 2016; Noguchi et al., 2007). Following affinity purification on a nickel-chelate column, the C-terminal thymosin-His_6_ moiety is preferentially cleaved off after the last, highly conserved Phe residue of actin by limited chymotryptic digestion, leaving actin with no foreign amino acids (Fig. 1A). Using this approach, we expressed and purified milligram quantities of fission yeast actin. The resulting G-actin was, however, incapable of polymerization following addition of KCl and Mg^2+^, as no filament formation was detected by high-speed centrifugation (Fig. 1B, left panel). Modulating the conditions for chymotryptic proteolysis did not produce polymerization-competent actin monomer. We postulated that the Tyr371 residue in close proximity to Phe375 (Fig. 1A) might be preferentially cleaved by chymotrypsin. Modifications at the actin C-terminus by either chemical crosslinking or proteolysis are known to reduce or abolish actin polymerization (Mossakowska et al., 1993; Otterbein et al., 2001; Strzelecka-Gołaszewska et al., 1995), and thus an alternative cleavage site could explain the inability of the actin to assemble. We thus substituted Y371 with a His residue, a non-aromatic amino acid found at this position in actin from many other species. Once this modified actin was expressed and purified, approximately twice the amount of chymotrypsin was required to cleave the tag from the actin(Y371H)-thymosin-β4-His_6_ variant compared with WT fusion protein, suggesting that F375 was a less favorable site for chymotryptic cleavage (Fig. 1C). The higher concentration of chymotrypsin introduced minor species due to secondary proteolysis (Fig. 1C, asterisks). More importantly, the purified actin(Y371H) showed a majority of the actin in the pellet following high-speed centrifugation, suggestive of filament formation (Fig. 1B, right panel)

**Figure 1.**
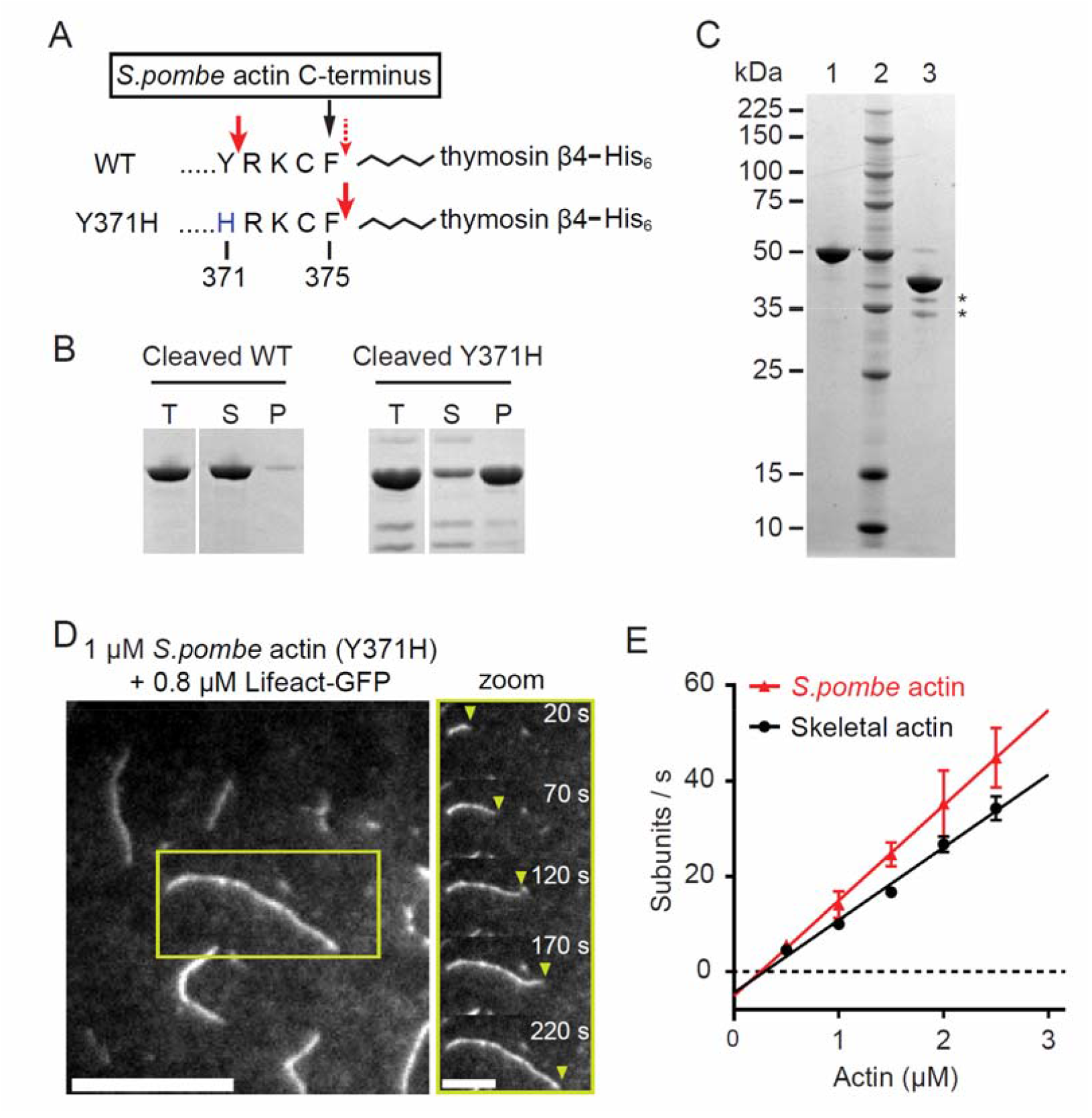
Expression and purification of polymerization-competent fission yeast actin from *Sf*9 insect cells. **(A)** Schematic illustration of the actin expression construct showing the C-terminal region of *S. pombe* actin (amino acid 371-375), followed by a linker (zigzag line), thymosin β4, and His_6_. The numbering of the amino acids takes into account that Met1 is not removed in yeast actins (Cook et al., 1991). The red bold arrows indicate the preferred chymotryptic cleavage sites in both constructs, the dotted arrow indicates the less preferred cleavage site in the WT construct. **(B)** 3 µM *S. pombe* actin, obtained from chymotrypsin-cleavage of the WT actin-thymosin β4 construct, was incubated in polymerization buffer (10 mM imidazole pH 7.5, 50 mM KCl, 4 mM MgCl_2_, 1 mM EGTA, 1 mM DTT, and 0.2 mM MgATP) at 30°C for 3 h. The Coomassie-stained gel shows the total (T), supernatant (S), and pellet (P) fractions following high-speed centrifugation. **(C)** Coomassie-stained 4-12% SDS-PAGE of purified *S. pombe* actin(Y371H)– thymosin β4-His_6_ before (lane 1) and after (lane 3) cleavage and removal of the thymosin β4-His_6_ moiety. Lane 2, molecular mass standards. * indicates minor bands resulting from proteolysis at sites other than F375. **(D)** Left, A representative image showing polymerized *S. pombe* F-actin (Y371H) filaments observed by TIRF microscopy. Scale bar, 10 µm. Right, a time montage showing an elongating *S. pombe* actin filament of the zoomed region (yellow open box) from the left panel. The yellow arrowheads denote the barbed end. Scale bar, 5 µm. (**E)** Actin polymerization rates of chicken skeletal actin (black circles) and *S. pombe* actin(Y371H) (red triangles) as a function of actin concentration, as observed by TIRF microscopy using LifeAct-GFP. The assembly and disassembly rate constants and critical concentrations obtained from the linear fits to the data are given in Table 1. Buffer: 10 mM imidazole pH 7.5, 50 mM KCl, 4 mM MgCl_2_, 1 mM EGTA, 1 mM DTT, 0.2 mM MgATP. Temp: 25 °C. n = 25-32 filaments per condition. Two independent actin preparations were used at each actin concentration.

### Polymerization of expressed fission yeast actin(Y371H)

The polymerization rates of expressed yeast actin(Y371H) and tissue-purified skeletal actin were quantified using total internal reflection fluorescence (TIRF) microscopy to visualize filament growth in real time. Polymerized F-actin was visualized with 0.8 µM Lifeact-GFP, a probe that binds F-actin with low affinity (K_d_ ∼2.2-10 µM) and which has been previously used to follow actin polymerization in real time (Courtemanche et al., 2016; Lu et al., 2015; Lu et al., 2016; Riedl et al., 2008). The expressed *S. pombe* actin(Y371H) was fully capable of polymerization (Fig. 1D), with a slightly faster polymerization rate constant (slope) than skeletal actin (19.9 vs 15.2 subunits s^-1^·µM^-1^) (Fig. 1E, Table 1). The depolymerization rate constants (y-intercept) of fission yeast actin and skeletal actin were similar (5.1 vs 4.4 subunits s^-1^·µM^-1^), as was the critical concentration (x-intercept) (0.25 µM vs 0.29 µM). Overall, the *in vitro* polymerization of expressed *S. pombe* actin(Y371H) resembled that observed with WT actin purified from fission yeast (Takaine and Mabuchi, 2007; Ti and Pollard, 2011) as well as skeletal actin.

**Table 1.** Polymerization rate constants.

| Actin species | Acetylated Tpm (Cdc8) | Assembly rate constant (subunits $\mu\text{M}^{-1}\cdot\text{s}^{-1}$) | Disassembly rate constant (subunits $\text{s}^{-1}$) | Critical concentration ($\mu\text{M}$) |
| --- | --- | --- | --- | --- |
| *S. pombe*<sup>a</sup> | - | 19.9 ± 0.4 | 5.1 ± 0.7 | 0.25 |
|  | + | 13.2 ± 1.3 | 1.6 ± 2.1 | 0.13 |
| skeletal | - | 15.2 ± 0.9 | 4.4 ± 1.6 | 0.29 |

<sup>a</sup>*S. pombe* actin (Y371H).

### *In vivo* function of the Y371H *S. pombe* actin variant

To assess whether *S. pombe* actin(Y371H) is fully functional *in vivo*, the variant allele was integrated into the fission yeast genome to replace the WT allele (Table S1). Lifeact-GFP was expressed in both WT and the actin(Y371H) variant strains to visualize the major actin structures. The yeast cells expressing actin(Y371H) properly form the contractile ring, have actin patches and actin cables that are indistinguishable from the WT cells, and have a comparable growth rate as WT cells (Fig. 2A-B). Additionally, the actin(Y371H) yeast cells have no discernable cell division defect, given the similar nuclei numbers per cell and the septum morphology in comparison to the WT cells (Fig. 2C, Table 2). Because actin(Y371H) supports fission yeast cytoskeleton function and exhibits WT-like activity, we used this variant for the rest of this study. For simplicity, actin(Y371H) is hereafter referred to as actin.

**Figure 2.**
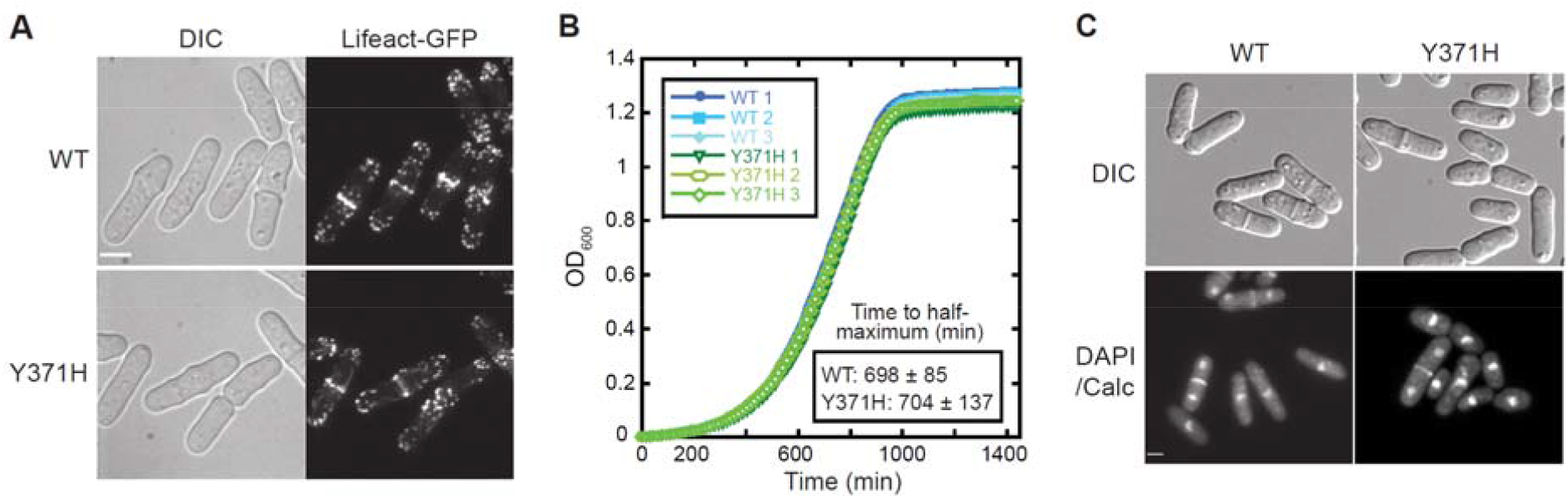
Growth characteristics of actin mutant Y371H fission yeast are similar to WT. **(A)** Micrographs of WT and actin(Y371H) fission yeast cells showing morphology (DIC) and organization of actin networks (Lifeact-GFP). Scale bar, 5 µm. **(B)** Growth curves of WT and actin(Y371H) strains. 3 replicates of each strain are shown. Average time to half-maximum OD_600_ of each strain was calculated from 15 replicates each. **(C)** Micrographs of WT and actin-Y371H cells in DIC (morphology), and methanol fixed cells stained with DAPI (nuclei) and Calcofluor (septa). Scale bar, 5 µm.

**Table 2.** Cell division characteristics of fission yeast strains expressing WT or Y371H actin.

|  | **WT** | **Y371H<sup>a</sup>** |
| --- | --- | --- |
|  | % of cells<sup>b</sup> |  |
| **One nucleus** | 77.3 ± 1.7 | 78.6 ± 1.8 |
| **Two nuclei** | 22.7 ± 1.7 | 20.8 ± 1.9 |
| **> two nuclei** | 0 | 0.68 ± 0.06 |
| **Normal septa** | 100 | 100 |
| **Abnormal septa<sup>b</sup>** | 0 | 0 |

<sup>a</sup> The strains used in this study are listed in Supporting information Table S1.
<sup>b</sup> Two replicates were analyzed for each strain. N >300 for each replicate.
<sup>c</sup> Abnormal septa are misplaced, unusually broad, or misshapen.

### Effect of fission yeast tropomyosin acetylation on binding to S. pombe actin

We compared how Tpm acetylation affects its binding to *S. pombe* F-actin. N-terminally acetylated fission yeast Tpm (Cdc8) was generated by co-expression of fission yeast N-α-acetyltranferase B complex with Tpm in bacteria (Johnson et al., 2010), because the acetyl-mimic (Ala-Ser) fission yeast Tpm was shown to have ∼10-fold lower affinity than the native acetylated isoform (Skoumpla et al., 2007). The purified acetylated Tpm (AcTpm) migrates slower than unacetylated Tpm on a urea/glycerol-based gel (Trybus, 2000), and shows only a single band confirming that all molecules are acetylated (Fig. 3A). Binding of AcTpm and unacetylated Tpm to *S. pombe* F-actin was assessed by a co-sedimentation assay. AcTpm has ∼3-fold higher affinity for actin than the unacetylated Tpm (*K*_app_ of 2.1 vs 0.8 × 10^6^·M^-1^, Fig. 3B). Both acetylated and unacetylated Tpm bind to actin cooperatively (Hill coefficient 2.4 vs 4.3), but acetylation enables cooperative binding of *S. pombe* Tpm at lower (3-4 fold) concentration.

**Figure 3.**
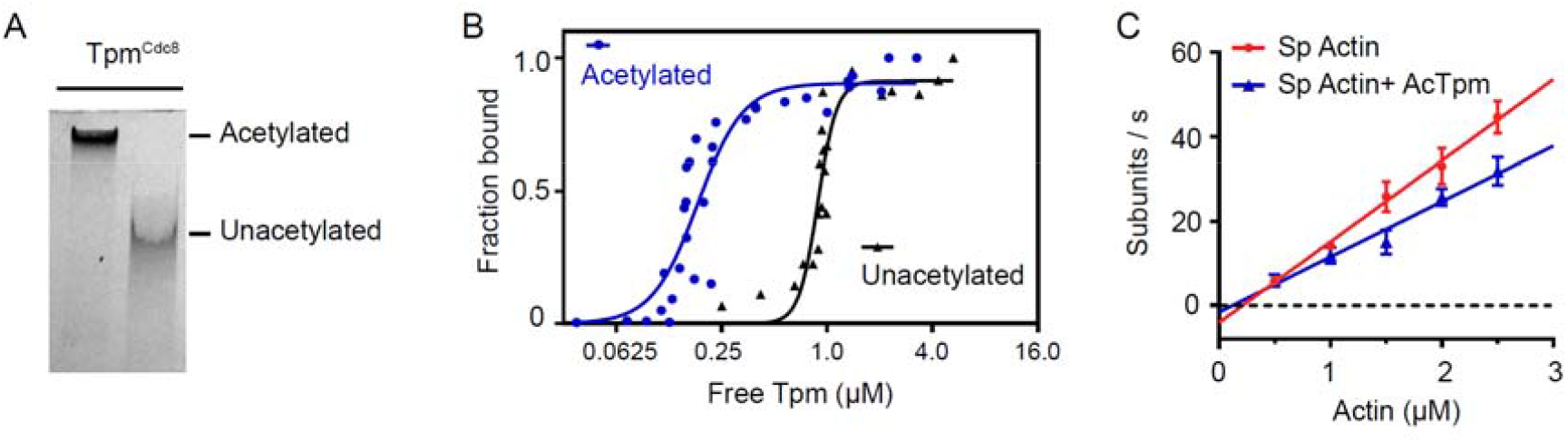
Acetylation of fission yeast Tpm increases the apparent affinity for fission yeast actin and slows actin polymerization dynamics. **(A)** Acetylated Tpm migrates slower than unacetylated Tpm on a charge gel. **(B)** Affinity of acetylated vs unacetylated *S. pombe* Tpm for *S. pombe* actin from a co-sedimentation experiment. Three experiments with two independent actin preparations were performed. Data points were fit to the Hill equation. *K*_app_ is 2.1 × 10^6^ M^-1^ for acetylated Tpm, and 0.8 × 10^6^ M^-1^ for unacetylated Tpm. The Hill coefficient is 2.4 for acetylated Tpm, and 4.3 for unacetylated Tpm. Buffer: 10 mM imidazole, pH 7.5, 150 mM NaCl, 2 mM MgCl_2_, 1 mM EGTA and 1 mM DTT. (**C)** *S. pombe* actin polymerization rates in the absence or presence of acetylated Tpm (AcTpm) observed by TIRF microscopy. The critical concentrations (0.22 µM for actin, 0.13 µM for actin-AcTpm) and the depolymerization rates (4.1 subunits s^-1^ for actin, 1.6 subunits s^-1^ for actin-AcTpm) were extrapolated from the linear fits. n = 22-39 filaments per condition. Two independent actin preparations were used for each actin concentration per condition.

Eighty percent of the Tpm in fission yeast is acetylated (Skoumpla et al., 2007), and thus it is of interest to measure the rates of *S. pombe* actin polymerization in the presence or absence of acetylated Tpm. Acetylated Tpm decreased the rate constants for both polymerization (19.2 vs 13.2 subunits s^-1^·µM^-1^) and depolymerization (4.1 vs 1.6 subunits s^-1^), leaving the critical concentration for actin assembly similarly low (0.25 µM vs 0.13 µM for AcTpm) (Fig. 3C, Table 1). These observations show that AcTpm slows *S. pombe* actin dynamics at both ends of the actin filament.

### Effect of tropomyosin on the interaction of fission yeast formin Cdc12 with S. pombe actin

We tested *in vitro* the hypothesis that fission yeast formins Cdc12 and For3 play a key role in specifying whether acetylated or unacetylated Tpm associates with the *S. pombe* actin filaments that they polymerize (Johnson et al., 2014). Specifically, does Cdc12 prefer acetylated Tpm and For3 unacetylated Tpm? We employed truncated, constitutively active fragments of both formins (FH1-FH2) in our *in vitro* studies for two major reasons: (1) full length diaphanous-related formins, such as Cdc12 and For3, are autoinhibited by DID-DAD interactions (Goode and Eck, 2007; Schönichen and Geyer, 2010), and (2) any interactions between formin and actin-Tpm copolymers are likely to occur through the FH2 domain that binds the filaments, while the N-terminal domains do not. To visualize how formins interact with Tpm, Cdc12 and acetylated Tpm(D142C) (Fig. 4A) were fluorescently labeled for visualization by TIRF microscopy. The D142C mutation in fission yeast Tpm was previously shown to be well tolerated *in vivo*, and *in vitro* exhibits an affinity for skeletal actin similar to WT-Tpm (Christensen et al., 2017). The *S. pombe* actin filaments were visualized by Lifeact-GFP. At Tpm concentrations ensuring complete binding to actin (≥1 Tpm:4 actin monomers), Tpm bound to actin in a highly cooperative fashion, with polymerizing actin filaments instantaneously decorated with Tpm. To temporally resolve the binding of Tpm to actin, the acetylated Tpm concentration was decreased (∼1 Tpm : 4 actin monomers). Across multiple experiments, we observed that while *S. pombe* actin filaments were being assembled by Cdc12, the acetylated Tpm did not first appear at the barbed end near the formin, but rather at various positions at a distance from the barbed end (Fig. 4B). This is similar to how acetyl-mimic *S. pombe* Tpm binds to growing skeletal F-actin filaments in the absence of formin (Christensen et al., 2017). As polymerization proceeded, most of the actin filaments eventually bound Tpm, but often left a gap next to the fast-growing actin barbed end (Fig. 4B). This observation argues against a direct and sustained interaction between formin Cdc12(FH1-FH2) domains and acetylated Tpm. When the same concentration of unacetylated Tpm was used, no Tpm binding on Cdc12 assembled filaments was observed (data not shown). Thus the higher affinity of acetylated Tpm for actin appears to be sufficient to account for the preference of acetylated Tpm binding to Cdc12 assembled networks.

**Fig 4.**
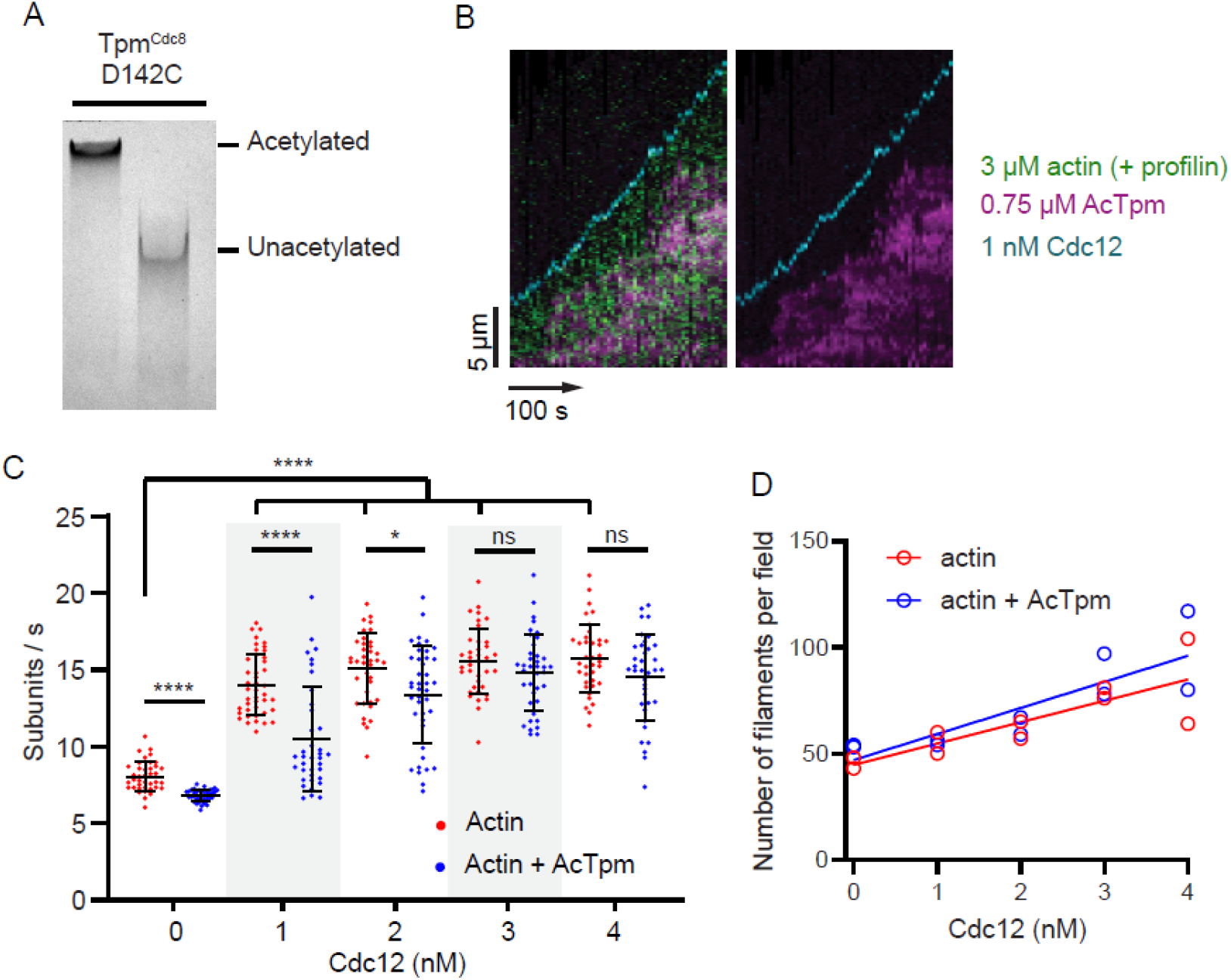
Acetylated fission yeast Tpm does not show a binding preference toward the fission yeast actin barbed end that is associated with contractile ring formin Cdc12 (FH1-FH2). **(A)** Acetylated Tpm Cdc8 (D142C) migrates slower than unacetylated Tpm on a charge gel. **(B)** Representative kymographs of an *S. pombe* actin filament barbed end elongation over time observed through TIRF microscopy in the presence of 3 µM *S. pombe* actin, 1 nM *S. pombe* formin Cdc12, 3 µM *S. pombe* profilin Cdc3, and 0.75 µM acetylated *S. pombe* Tpm (D142C). The formin (SNAP-Surface-649-labeled; cyan) and Tpm D142C (TMR-labeled; magenta) are directly labeled. Actin (green) is visualized using Lifeact-GFP. Left panel, 3-color composite. Right panel, 2-color composite of the left panel (only the formin and Tpm are shown). **(C)** Actin (1 µM) barbed end polymerization rates in the presence of 0-4 nM of formin Cdc12 and 1 µM of profilin, with (blue) or without (red) 1 µM acetylated Tpm. Actin was visualized with Lifeact-GFP. **(D)** The average number of total actin filaments observed in a 54 × 54 µm^2^ area after 2 min observed in TIRF microscopy in conditions described in C). n = 37-41 filaments per condition. Two independent actin preparations were tested in each condition. Kolmogorov-Smirnov test was performed. * *P* < 0.05, **** *P* < 0.0001, ns, not significant.

Similar to what was observed with self-assembled actin, acetylated Tpm slowed the rate of actin polymerized, but only at a low concentration of formin Cdc12 (1 nM) in the presence of profilin. At higher Cdc12 concentrations (2-4 nM), the rate of polymerization increased and diminished the slowing effect of Tpm (Fig. 4C, Table 3). The wider distribution of polymerization rates seen at the intermediate concentration range of Cdc12 (1-2 nM) may reflect a mixed population of actin filaments, some with and some without Cdc12 bound. Increasing Cdc12 concentration increased the number of filaments formed, but the presence of acetylated Tpm had little effect on filament number (Fig. 4D).

**Table 3.**
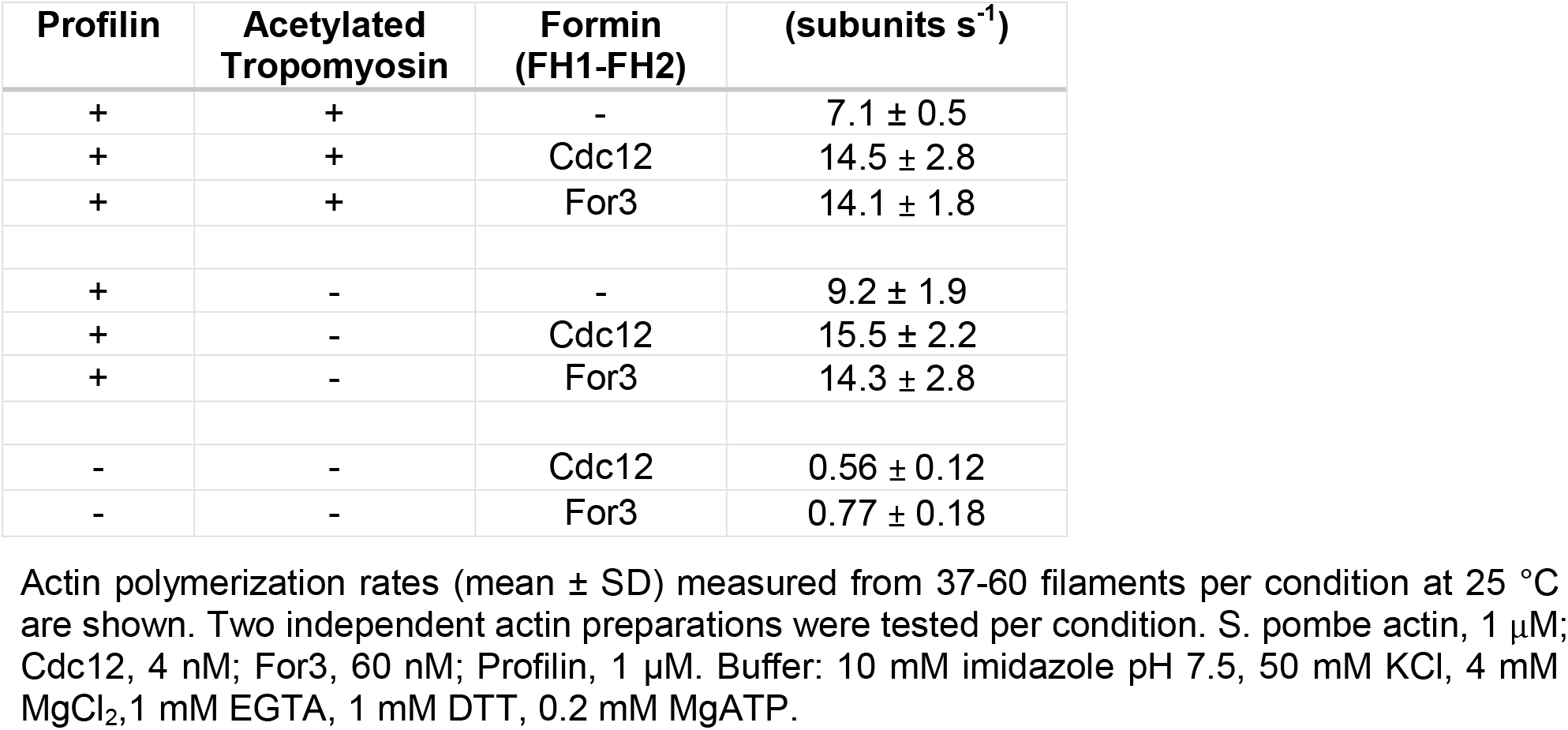
Effect of profilin, tropomyosin, and formin on the polymerization rate of *S. pombe* actin.

Cdc12 mediated skeletal actin filament elongation is gated by profilin (Kovar et al., 2003). The >10-fold slowing of polymerization observed when Cdc12 is added to *S. pombe* actin (Table 3) extends this finding and suggests that the capping of the actin barbed end by Cdc12 in the absence of profilin is an inherent characteristic of Cdc12.

### Effect of tropomyosin on the interaction of fission yeast formin For3 with S. pombe actin

We next tested whether unacetylated Tpm preferentially binds to For3 assembled *S. pombe* actin filaments. For3 expresses poorly in bacteria (Scott et al., 2011), and thus For3(FH1-FH2) was expressed using the baculovirus/insect cell system and high yields were obtained (5 mg/10^9^ cells, Fig. 5A). The concentrations of For3 (10-60 nM) required were, however, much higher compared to Cdc12, suggesting that it is less effective at polymerizing actin than Cdc12 (Fig. 5C).

**Fig. 5.**
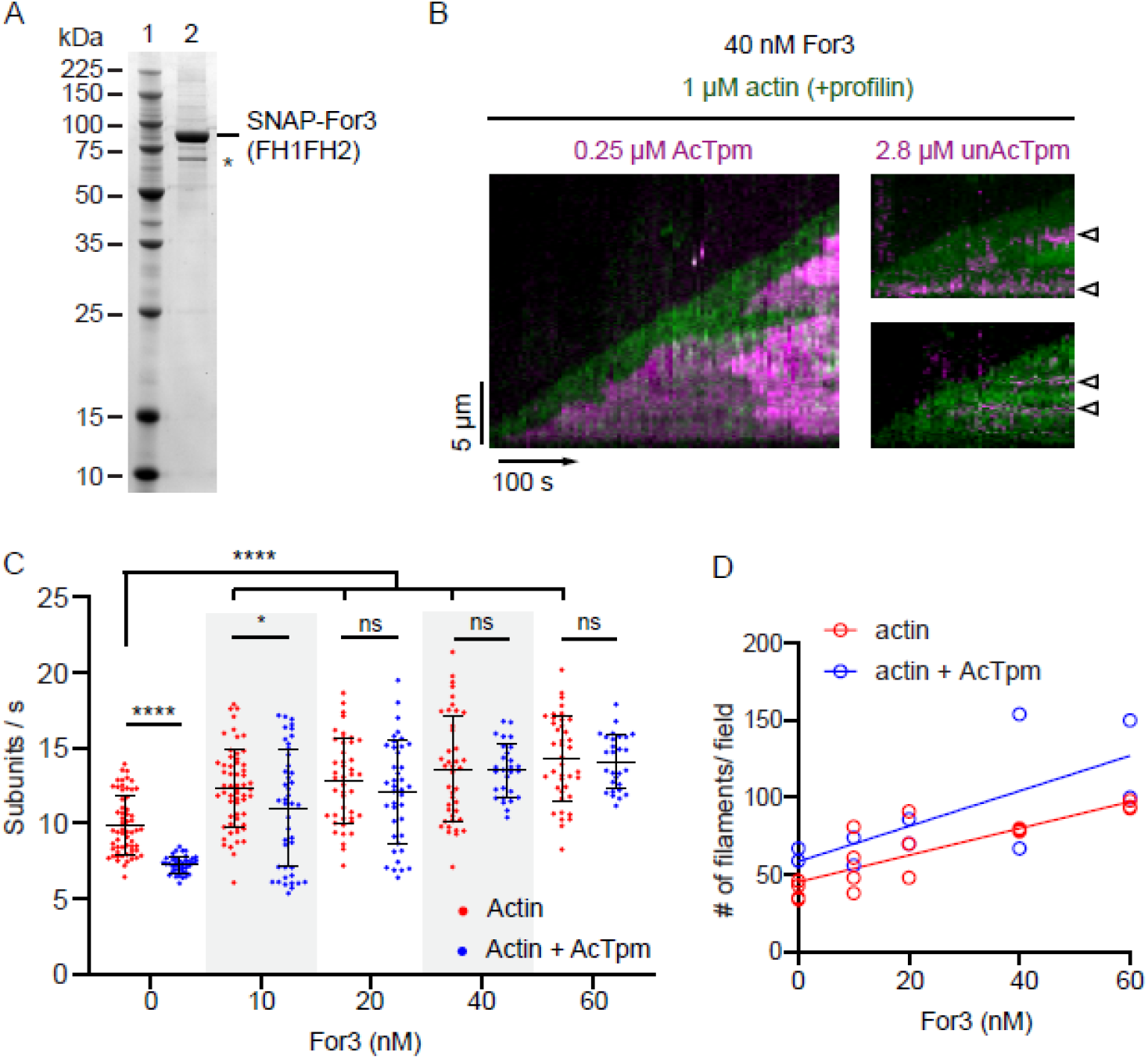
Fission yeast actin polymerized with cable formin For3 (FH1-FH2) recruits more acetylated than unacetylated Tpm and shows no preference for binding to the actin filament end. **(A)** 12 % SDS-PAGE showing SNAP-For3 (FH1-FH2)-HIS_6_ expressed and purified from *Sf*9 cells. Lane 1, protein standards. The * in lane 2 indicates a minor breakdown product that reacts with anti-HIS_6_ antibody. **(B)** Representative kymographs showing an *S. pombe* actin filament barbed end elongation in the presence of 1 µM *S. pombe* actin, 40 nM *S. pombe* For3, 1 µM *S. pombe* profilin Cdc3, and 0.25 or 2.8 µM *S. pombe* Tpm Cdc8 (D142C) over time. The actin (green) is visualized by using Lifeact-GFP, the Tpm D142C (TMR-labeled; magenta) is directly labeled. The For3 is unlabeled. Left panel: polymerizing actin is partially decorated by acetylated Tpm in the presence of 0.25 µM acetylated Tpm and 40 nM For3. Right panels, two examples showing very scarce regions of F-actin were bound to Tpm when 2.8 µM of unacetylated Tpm (D142C) and 40 nM For3 were present during actin polymerization, and no barbed end preference of Tpm binding to the actin filaments was observed. The open arrowheads showing the narrow, Tpm-bound regions on actin. **(C)** Actin (1 µM) barbed end elongation rates in the presence of 0-60 nM of formin For3 and 1 µM of profilin, with (blue triangle) or without (red circles) 1 µM acetylated Tpm. **(D)** The average number of total actin filaments observed in a 54 × 54 µm^2^ area after 2 min observed in TIRF microscopy in conditions described in C). n = 40-60 filaments per condition. Two independent actin preparations were tested in each condition. Kolmogorov-Smirnov test was performed. *\* P* < 0.05, **** *P* < 0.0001, ns, not significant.

As with Cdc12, TIRF microscopy was used to visualize binding of labeled Tpm(D142C) to actin in the presence of labeled For3. Acetylated Tpm bound to 1 µM *S. pombe* actin with higher affinity than unacetylated Tpm even in the presence of a high concentration of For3 (40 nM, Fig. 5B). A ∼10-fold higher concentration of unacetylated Tpm (2.8 vs 0.25 µM) was required for very limited binding to actin filaments. The short regions where unacetylated Tpm was bound on the actin filaments did not propagate (Fig. 5B, open arrow heads), and some regions eventually dissociated from actin filaments. This observation is consistent with the actin-Tpm co-sedimentation experiments in the absence of formin and profilin, which showed that the unacetylated Tpm reaches cooperative binding to actin only at very high Tpm concentration (Fig. 3B). Visually, neither acetylated nor unacetylated Tpm appeared to associate with the elongating barbed end (Fig. 5B).

For3, in the presence of profilin, increased the rate of polymerization of *S. pombe* actin (Fig. 5C). Similar to Cdc12, high concentrations of For3 diminishes the slowing effect of acetylated Tpm on actin polymerization (Fig. 5C). The number of actin filaments increases with For3 concentration, with slightly more filaments in the presence of acetylated Tpm (Fig. 5D). As previously reported using skeletal actin, in the absence of profilin, For3 caps the fission yeast *S. pombe* actin barbed end as observed by the >10-fold slowing of polymerization rate (Table 3) (Scott et al., 2011).

### Interaction of *S. pombe* actin with *S. pombe* Myo2 and Myo51

We compared the ability of full-length Myo2, a class II myosin that is essential for contractile ring assembly in fission yeast, to move skeletal versus *S. pombe* actin in an *in vitro* motility assay. NHSR-labeled *S. pombe* actin (9-10% labeled) was used because fission yeast actin has extremely low affinity for phalloidin (Ti and Pollard, 2011). We observed similar speed distributions whether *S. pombe* actin (Fig. 6A, red) or skeletal actin (Fig. 6A, black) was used, with mean values of 0.67 µm/s and 0.72 µm/s respectively (Fig. 6A, upper panel, Table 4). When acetylated tropomyosin was present, the Myo2 motility was reduced to 0.5 µm/s with both actin species (Fig. 6A, lower panel), consistent with observations in previous studies on Myo2, whether the myosin was isolated from fission yeast or *Sf*9 cells (Pollard et al., 2017; Stark et al., 2010).

**Fig 6.**
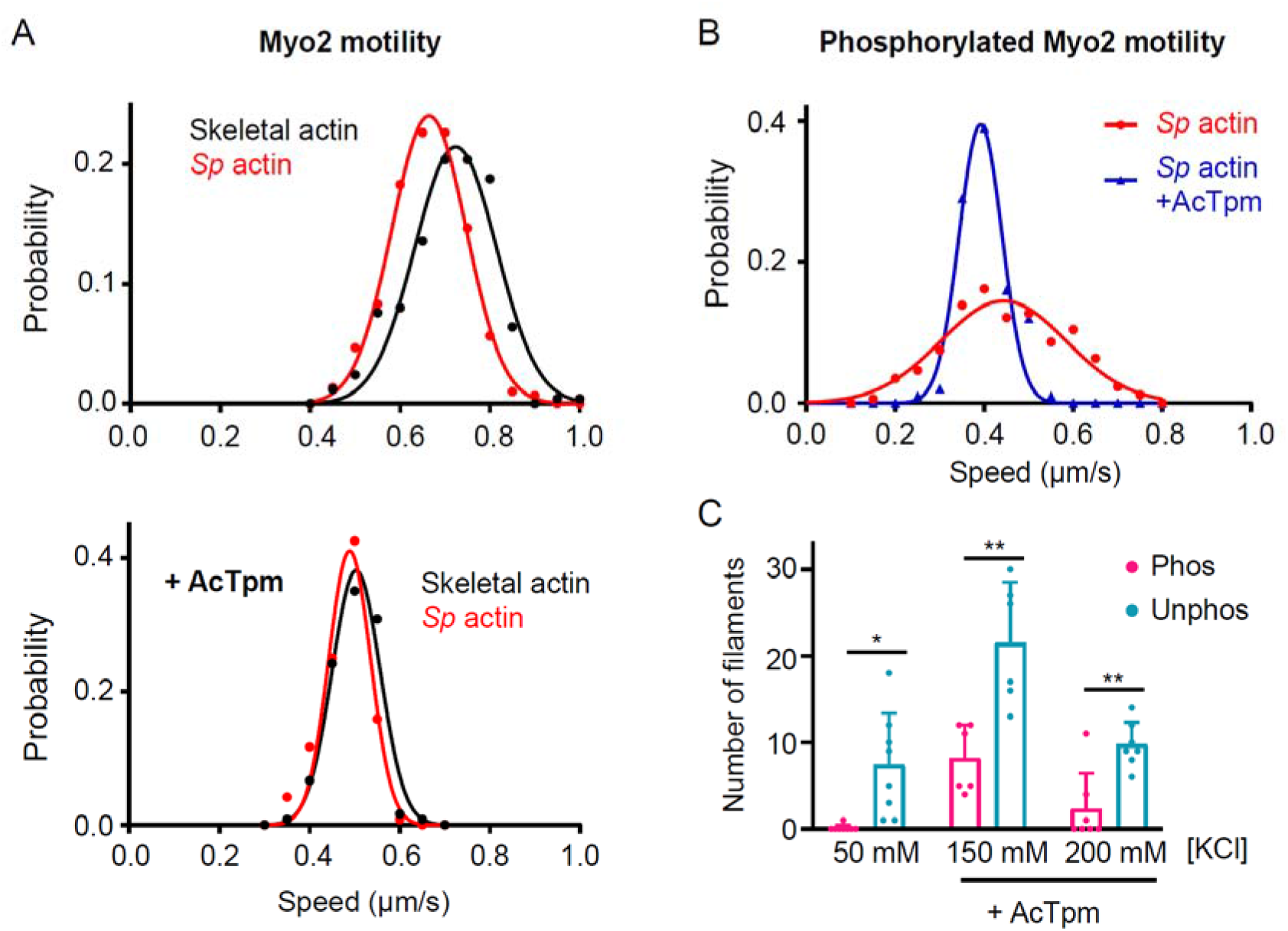
Myo2 moves fission yeast or skeletal actin filaments similarly in an *in vitro* motility assay. **(A)** *In vitro* motility speeds of unphosphorylated fission yeast contractile ring myosin Myo2 (full length), with directly labeled (NHS-rhodamine) skeletal actin (black) and *S. pombe* actin (red) at 30 °C in buffer containing 50 mM KCl, 0.5 % methylcellulose, and 1 mM MgATP. Lower panel, motility speed distribution in the presence of 2 µM acetylated Tpm. n = 120-300 filaments per condition for each actin. Statistics are shown in Table 4. (**B)** *In vitro* motility speeds of phosphorylated Myo2 with directly labeled *S. pombe* actin with (blue triangles) or without (red circles) acetylated Tpm. The bare actin shows a wider distribution of speeds compared to actin-Tpm. n = 170-300 filaments per condition. **(C)** Number of *S. pombe* actin filaments bound to the phosphorylated (magenta) or unphosphorylated (cyan) Myo2-coated surface (128 µm × 128 µm), in the absence of methylcellulose at different KCl concentrations. Acetylated Tpm was added to 2 µM when indicated. Note that Myo2 did not bind actin filaments at 150 mM KCl in the absence of Tpm (Pollard et al., 2017). The data were collected from experiments using two independent actin preparations. Unpaired *t*-test with Welch’s correction, * *P* < 0.05, ** *P* < 0.01.

**Table 4.** Actin *in vitro* motility speeds.

| myosin | conditions | AcTpm | *S. pombe* actin ($\mu\text{m s}^{-1}$) | Skeletal actin ($\mu\text{m s}^{-1}$) | Comparison between the two actins |
| --- | --- | --- | --- | --- | --- |
| **Myo2** | Unphosphorylated | - | $0.66 \pm 0.08$ | $0.72 \pm 0.09$ | n.s |
| | | + | $0.49 \pm 0.05$<br>**** | $0.50 \pm 0.05$<br>**** | n.s |
| | Phosphorylated | - | $0.44 \pm 0.14$ | nd | |
| | | + | $0.39 \pm 0.05$ * | nd | |
| **Myo51-Rng8/9** | 50mM KCl | - | $1.35 \pm 0.17$ | $1.39 \pm 0.14$ | n.s |
| | 100mM KCl | - | $1.62 \pm 0.18$ | $1.71 \pm 0.12$ | n.s |
| | 150mM KCl | - | $1.87 \pm 0.15$ | $1.81 \pm 0.16$ | n.s |
|  | 150mM KCl | + | 0 | 0 | n.s |

Motility speed (mean ± SD) tracked from 80-300 filaments per condition are shown. Two independent actin preparations were used per condition. Nd, not determined. Actin gliding motility assays were performed at 30 °C in buffer containing 25 mM imidazole pH 7.5, 50 mM KCl (or otherwise indicated in the table), 4 mM $\text{MgCl}_2$ , 1 mM EGTA, and 10 mM DTT, oxygen scavenger, 1mM MgATP, and 0.5 % (for Myo2) or 0.5-0.7 % (for Myo51) methylcellulose. 2 $\mu\text{M}$ acetylated Tpm was added when actin-Tpm was tested. Unpaired t-tests were performed to compare motility speeds on *S. pombe* versus skeletal actin (right most column). n.s, not significant. Unpaired t-tests with Welch's correction (not assuming same SD) were performed to compare the motility speeds on bare versus AcTpm-bound actin, indicated next to the motility speed. \* $P < 0.05$ , \*\*\*\* $P < 0.0001$ .

We then tested *S. pombe* actin motility using Myo2 that is phosphorylated by fission yeast Pak1 kinase (Loo and Balasubramanian, 2008; Pollard et al., 2017). Phosphorylated Myo2 has low affinity for actin and shows a broad speed distribution that is narrowed when Acetylated Tpm is present (Fig. 6B, Table 4), similar to what we previously observed using skeletal actin (Pollard et al., 2017). Using a modified motility assay to quantify actin filament binding, phosphorylation of Myo2 decreases its affinity for *S. pombe* actin whereas acetylated tropomyosin increases binding (Fig. 6C, Table 4). These results using fission yeast actin validated our recent characterization of Myo2 with skeletal muscle actin (Pollard et al., 2017).

Lastly, we examined the interaction between fission yeast actin with full-length Myo51-Rng8/9, a myosin complex that is recruited to both the contractile ring and the actin cables in a tail-dependent manner (Lo Presti et al., 2012; Wang et al., 2014). Here we found similar actin gliding speed distributions between skeletal and fission yeast actins at multiple ionic strengths (Figure 7A). Gliding speed increases with ionic strength, likely because Myo51 has a high duty ratio (Moore et al., 2001; Tang et al., 2016; Wang et al., 2000). Further, the presence of acetylated tropomyosin stopped gliding motility (Table 4) due to the tail binding of the Myo51 complex to the acetylated tropomyosin bound actin (Fig. 7B), as we reported previously using skeletal actin (Tang et al., 2016). Taken together, Myo51-Rng8/9 interacts with *S. pombe* actin in a similar manner as skeletal actin.

**Fig 7.**
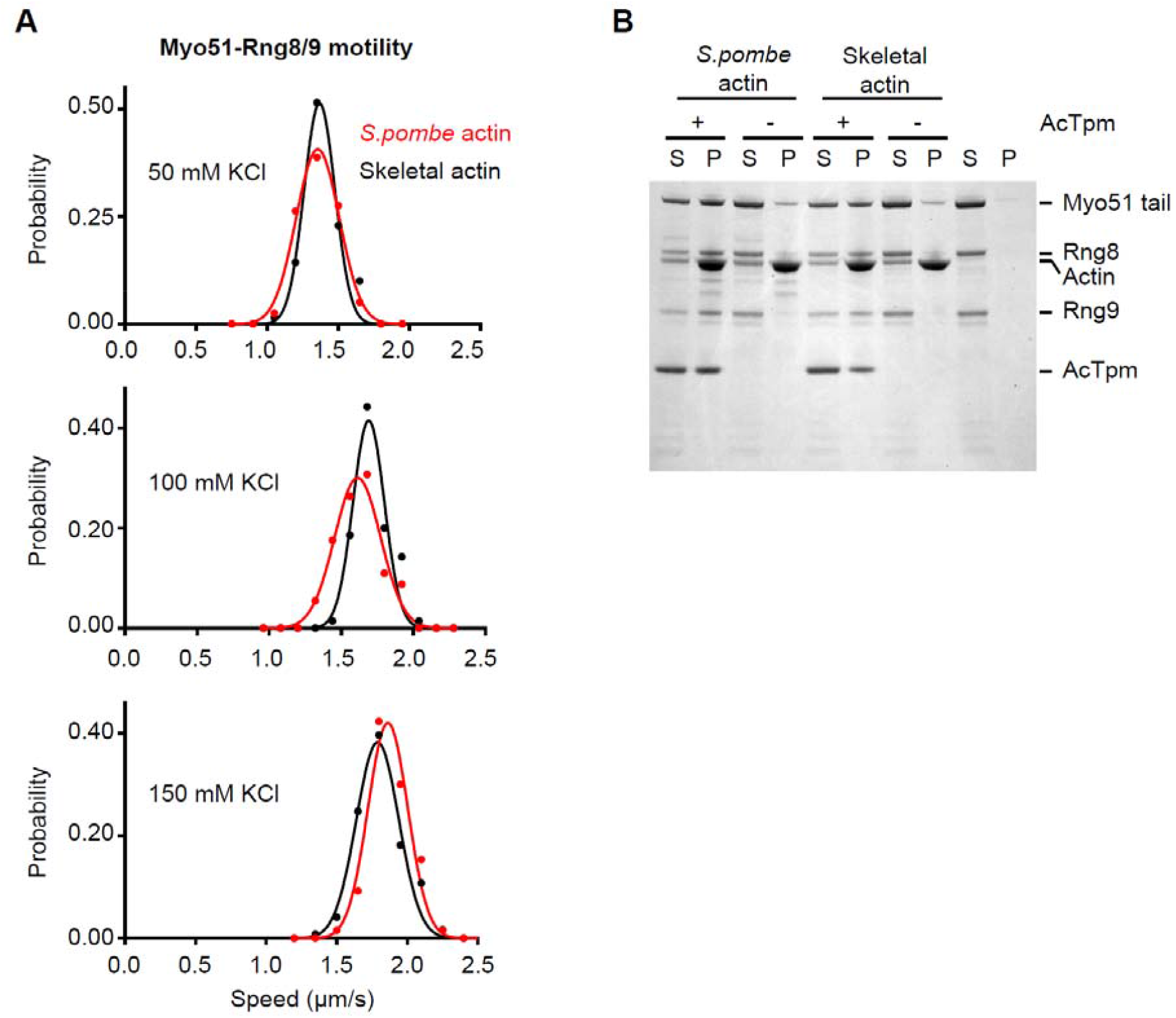
Myo51-Rng8/9 moves fission yeast or skeletal actin filaments similarly in an *in vitro* motility assay. **(A)** *In vitro* motility speeds of fission yeast contractile ring and actin cable myosin Myo51-Rng8/9 (full length) with directly labeled (NHS-rhodamine) skeletal actin (black) and *S. pombe* actin (red) at 30 °C, in the buffers containing 50-150 mM KCl, 0.5-0.7% methylcellulose, and 1.5 mM MgATP. n = 80-130 filaments per condition for each species. Data were collected from experiments using two independent actin preparations. **(B)** 1 µM of Myo51 tail-Rng8/9 pellets with either *S. pombe* or skeletal actin in the presence of acetylated Tpm. Actin concentration, 4 µM; acetylated Tpm, 2 µM; 150 mM NaCl. The supernatant (S) and pellet (P) fractions are resolved on a 12 % SDS-PAGE.

## Discussion

A long-standing question in the field is how tropomyosins are sorted to different actin structures within a cell. As a conserved family, tropomyosins are α-helical coiled-coil proteins that form head-to-tail interactions that bind actin filaments cooperatively. Previous studies have revealed that tropomyosin can either compete with or promote the association of actin-binding proteins with actin (Christensen et al., 2017; Christensen et al., 2019; Hsiao et al., 2015; Skau and Kovar, 2010; Winkelman et al., 2016). But very little is understood regarding how the different isoforms of Tpm (up to ∼40 isoforms in mammalian cells) ‘select’ their actin network. An attractive hypothesis that was proposed was that formins may influence F-actin structural conformation or directly interact with Tpm to pair the actin with a specific Tpm isoform. A few studies provided evidence on a potential direct interaction between Tpm and formin *in vitro* and *in vivo* (Alioto et al., 2016; Johnson et al., 2014; Wawro et al., 2007). By combining fission yeast actin and a minimum set of fission yeast proteins, here we examined whether formin determines if acetylated or unacetylated tropomyosin binds to the actin filament it assembles.

Our results do not support the ideas that the constitutively active fission yeast FH1-FH2 domains influence tropomyosin sorting. We found acetylated Tpm bound to actin filaments with much higher affinity under all conditions we tested, independent of the presence of formins. The unacetylated Tpm has a lower apparent affinity for actin and requires 4-fold higher concentration to reach cooperativity in comparison to acetylated Tpm. This is consistent with the established role of acetylation in stabilizing tropomyosin N-terminal α-helical conformation and hence the interaction between Tpm N- and C-terminals to allow cooperative binding to actin (Frye et al., 2010; Greenfield et al., 1994). In the context of fission yeast, these observations suggest that achieving cooperativity is the rate-limiting step for unacetylated Tpm to ‘out-compete’ the acetylated Tpm for F-actin association, especially given that the unacetylated Tpm accounts for only 20% of the total Tpm *in vivo* (Skoumpla et al., 2007). Indeed, it was also reported that in mammalian cells, the two major formins mDia1 and mDia3 are not essential for specifically recruiting different tropomyosin isoforms to stress fibers. Instead, the incorporation of tropomyosin isoforms was suggested to be influenced by their respective concentrations (Meiring et al., 2019). A recent study showed that different mammalian Tpm isoforms associate with actin filaments with different dynamics, and compete with other actin-binding proteins in an isoform-specific manner *in vitro* (Gateva et al., 2017). Further, tropomyosins contain post-translational modifications (Carman et al., 2021). In fission yeast, phosphorylation of S125 in Cdc8 was shown to significantly reduce its apparent actin binding affinity and lead to destabilization of contractile ring and actin cable formation (Palani et al., 2019). It should be noted that our formin constructs (For3 and Cdc12) only contain the portions that interact with actin and profilin-actin (FH1-FH2) and thus we did not directly test the possibility that the N-terminal domains that are involved in formin activation/autoinhibition and recruitment (the armadillo repeat regions that contain the DID, GTPase-interacting, and other regulatory regions) may somehow interact with Tpm. A main challenge is that the full-length formins are autoinhibited, requiring additional factors to activate them *in vitro*. Additionally, our reconstitution is based on the assumption that Tpm decorates the actin filament where the FH2 domain is known to directly interface. To date, there is no published evidence for direct interactions occurring between the formin N-terminus and actin-tropomyosin. We also did not test the effects of posttranslational modifications of the formins on differential Tpm interactions. Phosphorylation events have been shown to inhibit Cdc12 from its roles in contractile ring assembly (Bohnert et al., 2013; Willet et al., 2018), but no studies show direct phosphorylation of For3 (of 35 publications mentioning For3 listed on PubMed.gov). Until evidence to support such mechanisms arises, the possible roles of phosphorylation on formin-Tpm binding remain highly speculative. Overall, a formin-regulated Tpm isoform sorting mechanism may require additional molecular components (including formin domains outside of the FH1 and FH2), which have not been clearly identified.

Our study also describes a robust method for the purification of *S. pombe* actin in a heterologous system. We first attempted to express N-terminally tagged fission yeast actin (N-FLAG-TEV-actin) using the Sf9/baculovirus system. For this construct, Met was substituted for the Gly/Ser in the TEV recognition site, which allowed TEV cleavage while retaining only residues native to *S. pombe* actin, since Met1 is not cleaved off in yeast as it is in mammals (Cook et al., 1991). However, this method produced insufficient amounts for our intended biochemical study, possibly because this actin can polymerize in the *Sf*9 cells and dysregulate the cytoskeleton. Recombinant expression of an actin-thymosin fusion in eukaryotic cells overcomes the hurdle of expression, presumably by sequestering actin as a monomer and thus preventing negative effects of ectopic overexpression. This expression strategy was previously used to enable the mechanistic characterization of toxic actin mutants in *Dictyostelium* (Noguchi et al., 2007), and the disease-causing human vascular smooth muscle actin mutants (Lu et al., 2015; Lu et al., 2016). During the course of preparing our study for publication, a strategy similar to ours was employed in a study where the budding yeast *Pichia pastoris* was used as the expression host (Hatano et al., 2018). Their construct retained the native residue Y371, but chymotrypsin did not cleave at this residue while it did in our *Sf*9 cell expressed actin (see Fig. 1). We speculate that this could be due to how the thymosin was cleaved off when the actin was bound to the nickel resin in the Hatano et al. study, or perhaps the Tyr was post-translationally modified in *Pichia* and not in *Sf*9 cells, thus making it a less preferable substrate.

Our recent *in vitro* characterization of fission yeast myosins Myo2 and Myo51 revealed biochemical behaviors that are unprecedented in each of their myosin classes, and here we show that our key prior results obtained using skeletal actin are recapitulated using expressed *S. pombe* actin. We found that Myo2 motility speeds are similar between *S. pombe* and skeletal actins. Additionally, phosphorylation of Myo2 decreased its affinity for *S. pombe* actin, but the addition of the acetylated Tpm increased its affinity and narrowed its speed distribution as was previously observed with skeletal actin. Further, we showed here that Myo51-Rng8/9 tail can specifically attach to acetylated Tpm bound-*S. pombe* actin, but not actin alone, and that Myo51 motility speeds are identical on *S. pombe* and skeletal actin without Tpm. The similar biochemical characteristics of two fission yeast myosins with either fission yeast or skeletal actin shown here further supports the proposed functions of these myosins based on studies using skeletal actin. The conservation of *S. pombe* myosin activity on skeletal and *S. pombe* actins is consistent with a recent cryo-EM study showing that the myosin footprint on actin is conserved between even highly divergent species (Robert-Paganin et al., 2021).

Other studies, however, have shown that myosin kinetics can vary with actin isoform (Cook et al., 1993; Stark et al., 2011; Takaine and Mabuchi, 2007). For example, it has been suggested that negative charges at the N-terminus of the actin subunit can affect how actin filaments interact with the *S. cerevisiae* myosin motor domain (Stark et al., 2011). Given that we found no significant differences between muscle and *S. pombe* actin in motility assays employing Myo2 and Myo51, there may be key structural differences in how these two yeasts have diverged over ∼400 million years since their common ancestor (Sipiczki, 2000). Work from the Manstein laboratory has suggested that skeletal-α and cytoplasmic -β and -γ actins can have differential impacts on the kinetics of mammalian non-muscle myosin-II isoforms but not on Myosin-IC (Müller et al., 2013; Reindl et al., 2022), supporting our findings that some myosins are insensitive to actin isoform differences. Additionally, Tpm isoforms further differentially fine-tune class-specific myosin motor activities in their associated structures in mammalian cells (Reindl et al., 2022). Intriguingly, recent work has suggested that sequence differences in actin ortholog at the DNA/gene level can have a greater impact on growth and survival of *S. cerevisiae* mutants than the protein sequence *per se* (Boiero Sanders et al., 2022). *S. cerevisiae* expressing human β-actin from the endogenous actin locus using *S. cerevisiae* codon usage survive but are deficient in growth and polarity, and the actin patches in these mutants are resistant to CK-666, a small molecule inhibitor of Arp2/3 (Boiero Sanders et al., 2022). These findings suggest that human-β and *S. cerevisiae* actins exhibit biases towards binding different sets of *S. cerevisiae* actin binding proteins. Thus, the rules governing compatibility between proteins from heterologous systems are undefined, underscoring the need to perform biochemical experiments in a completely cognate system whenever possible.

The results reported here show that some myosins can tolerate different actin isoforms as they interact through a conserved structural interface (Robert-Paganin et al., 2021), yet the rules for actin isoform compatibility are unclear for many other actin binding proteins. A few mammalian tropomyosin isoforms can complement fission yeast Tpm deficient strains (Balasubramanian et al., 1992; Clayton et al., 2014; Skoumpla et al., 2007). However, there are considerable differences between fission yeast actin and skeletal actin. Two previous studies reported that purified fission yeast actin differs in its *de novo* polymerization kinetics, as well as its interactions with fission yeast profilin, cofilin, and Arp2/3 in comparison to skeletal actin (Takaine and Mabuchi, 2007; Ti and Pollard, 2011). For example, fission yeast profilin does not stimulate nucleotide exchange on fission yeast actin monomer but does so on skeletal actin (Ti and Pollard, 2011). In addition, *S. pombe* actin binds to phalloidin poorly, which indicates a substantial difference in filament thermal fluctuation rates and/or the phalloidin binding surface comparing to skeletal actin (De La Cruz and Pollard, 1996; Ti and Pollard, 2011). Altogether, resolving the species/isoform-specific characteristics of actin and actin binding proteins will require reconstitution of additional components with native protein sequences and even faithful post-translational modifications.

## Material and Methods

### Sf9/baculovirus constructs

Human thymosin β4 sequence was linked C-terminally to S*f*9 codon-optimized (Integrated DNA Technologies) fission yeast actin cDNAs (WT and Y371H) bridged by a 14 aa-linker (ASSGGSGSGGSGGA), and ligated into pFastbac1 vector (ThermoFisher). The Met1 of fission yeast actin was kept in both sequences because yeast actin N-termini are not cleaved (Cook et al., 1991). The cDNA of fission yeast formin For3 FH1-FH2 domain (amino acid 718-1265) was fused to an N-terminal SNAP tag sequence and a C-terminal HIS_6_ tag, and cloned into pFastbac1 vector (ThermoFisher). A previously published N-terminally SNAP-tagged Cdc12 FH1-FH2 His_6_ (Cdc12 amino acid 882-1390) was used (Kovar et al., 2003; Scott et al., 2011; Zimmermann et al., 2017). Fission yeast tropomyosin (Cdc8) cDNAs (WT and D142C) were ligated into pET3a vector (Novagen).

### Actin mutagenesis and S. pombe strains

The fission yeast strains used in this work are listed in Supplementary Table S1. To generate the Y371H actin strain, the WT actin gene was amplified from WT fission yeast genome and ligated into a pFA6a-KanMX6 vector using SalI and PacI restriction sites. The Y371H mutation was introduced using QuikChange Site-Directed Mutagenesis kit (Agilent Technologies, Inc., Santa Clara, CA), and the actin mutant-Kan cassette was PCR-amplified and transformed into WT fission yeast via lithium acetate transformation. The Y371H mutation integrated into the genome was confirmed by sequencing. The resulting cassette integration replaced the endogenous terminator with the TEF terminator and integrated the kanMX6 selection gene downstream of actin before 1476148 bp on chromosome II. Actin Y371H strain expressing Lifeact-GFP was constructed through a genetic cross on SPA5S plates followed by tetrad dissection on a YE5S plate.

### Fluorescence microscopy and imaging conditions for visualizing fission yeast

For live cell imaging, cells were grown overnight in YE5S (yeast extract + five supplements) at 25° C and subcultured in EMM5S (Edinburgh minimal medium + five supplements) for 20 hours prior to imaging. For visualization of Lifeact-GFP, cells were imaged directly on a glass slide. Z-stacks of 13 slices were acquired at 300 ms exposure on a Zeiss Axiovert 200M fitted with a Cascade 512B EM-CCD camera (Photometrics, Tuscon, AZ) and equipped with a Yokogawa CSU-10 spinning disk unit (McBain, Simi Valley, CA) and a 100×, 1.4 NA objective illuminated with a 50 mW 473 nm DPSS laser controlled by Metamorph software (Molecular Devices, Sunnyvale, CA).

To stain cells with DAPI and Calcofluor, cells were grown in YE5S at 25 °C for 24-36 h, and fixed in 100 % methanol chilled to 4 ºC. The fixed cells were incubated with 3.3 µg/mL Calcofluor (Fluka Analytical, Sigma Aldrich, St. Louis, MO) in 50 mM sodium citrate at 37 °C for 1 min, washed with 50 mM sodium citrate, and resuspended in 50 mM sodium citrate containing 6.5 µg/mL DAPI (Life Technologies, Carlsbad, CA). The stained cells were kept on ice prior to imaging as previously described (Christensen, et al. 2017).

### Cell growth measurement by optical density

To obtain the cell growth curve over time, fission yeast were cultured in YE5S broth at 25°C for 24 h, then diluted to three different OD_600_ (0.06, 0.03, and 0.015) each in 200 µl of YE5S liquid media in a 96-well, flat bottom, clear polystyrene plate (Corning Incorporated, Corning, NY). The plate was shaken with a 4 mm orbital amplitude and the OD_600_ readings were recorded every 15 min for 24 h at 25-30 °C using a Tecan Infinite 200 PRO plate reader (Tecan, Durham, NC).

### Protein purification

The *Sf*9 cells expressing the actin-thymosin β4-HIS_6_ fusion proteins were pelleted and resuspended in HIS-binding buffer (10 mM HEPES pH 7.5, 300 mM NaCl, 0.25 mM CaCl_2_, 0.5 mM MgATP, 7 mM β-mercaptoethanol, 0.5 mM 4-(2-Aminomethyl) benzenesulfonyl fluoride hydrochloride (AEBSF), 5 mM benzamidine, 0.5 mM phenylmethylsulfonyl fluoride (PMSF), 5µg/mL leupeptin) on ice, and sonicated. The cell debris was spun down at 250,000 g. The supernatant was then incubated with HIS-Select nickel affinity gel (Sigma-Aldrich) for 1 h. The resin was washed with HIS-binding buffer and wash buffer (10 mM imidazole, 10 mM HEPES pH 7.5, 300 mM NaCl, 0.25 mM CaCl_2_, 0.25 mM MgATP, 7 mM β-mercaptoethanol), the fusion protein was eluted with elution buffer (200mM Imidazole, 10mM HEPES pH7.5, 300 mM NaCl, 0.25 mM CaCl_2_, 0.25 mM Na_2_ATP, 7 mM β-mercaptoethanol), and dialyzed against G-buffer (5 mM Tris-HCl pH 8.26 at 4°C, 0.2 mM CaCl_2_, 0.1 mM NaN_3_, 2 mM DTT, 0.2 mM Na_2_ATP) for 3 d and clarified by centrifugation at 300,000 g for 45 min. To obtain untagged actin, the fusion protein was incubated with chymotrypsin (weight ratio of chymotrypsin/actin is 1:66 for Y371H, and 1: 125 for WT) for 15 min at 23°C and the reaction was stopped by adding 2 mM PMSF and 2.5 mM AEBSF. The mixture was clarified by centrifugation at 300,000 g for 15 min, and the cleaved actin and thymosin β4-His_6_ moiety were separated by ion-exchange chromatography with SuperQ-5PW resin (Tosoh Bioscience, Japan) using a 0-0.5 M NaCl gradient in G-Buffer. The appropriate actin fractions were pooled and dialyzed against G-buffer for 3 d, clarified by centrifugation, and concentrated by Amicon-Ultra filtration (EMD Millipore).

The *Sf*9 cells expressing SNAP-For3 (FH1-FH2)-HIS_6_ was pelleted and sonicated in lysis buffer (50 mM Na_2_PO_4_, pH 8, 500 mM NaCl, 10 % (vol/vol) glycerol, 10 mM imidazole, 10 mM β-mercaptoethanol, 1 µg/mL leupeptin, 0.5 mM AEBSF, 5 mM benzamidine, 0.5 mM PMSF. The lysate was clarified by centrifugation at 250,000 g for 30 min and the supernatant was incubated with HIS-Select nickel affinity gel (Sigma-Aldrich) for 1 h at 4 °C. The bound protein was washed with lysis buffer and eluted with elution buffer (50 mM Na_2_PO_4_, pH8, 250 mM imidazole, 500 mM NaCl, 10 % (vol/vol) glycerol, and 10 mM β-mercaptoethanol). The fractions were pooled and concentrated by Amicon-Ultra filtration (EMD Millipore), clarified by centrifugation at 300,000 g for 30 min, dialyzed against the storage buffer (10 mM imidazole pH 7.4, 1 mM EDTA, 500 mM NaCl, 50 % (vol/vol) glycerol, and 2 mM DTT), and stored at -20 °C, or flash frozen and stored at -80 °C. The For3 was diluted and spun at 400,000 g for 30 min each time before use and the protein concentration in the supernatant was determined by Bradford protein assay (ThermoFisher). The protein gradually loses its activity on ice overnight.

The Myo2-C-biotin-FLAG, Myo51-Rng8/9-C-biotin-FLAG and N-FLAG-biotin-Myo51-Tail-Rng8/9 were expressed in Sf9/baculovirus system and purified as previously described (Pollard et al., 2017; Tang et al., 2016). Briefly, *Sf*9 cells co-infected with baculovirus encoding the components of each myosin complex were harvested after 72 h, pelleted and resuspended in ice-cold myosin lysis buffer (10 mM imidazole pH 7.5, 300 mM NaCl, 5 mM MgCl_2_, 1 mM EGTA) supplemented with 2 mM DTT, 1 µg/mL leupeptin, 0.5 mM AEBSF, 5 mM benzamidine, and 0.5 mM PMSF. For full length Myo51-Rng8/9, the myosin lysis buffer was supplemented with 20 µg/mL of bacterially expressed light chains (Cam1 and Cdc4). Cells were lysed by sonication and the cell debris was removed by centrifugation in the presence of 2 mM MgATP. The supernatants were incubated with FLAG affinity resin for 1 h, the resin was washed in lysis buffer without supplements, and the bound proteins were eluted after using FLAG peptide at 100 µg/mL. The eluate was concentrated using Amicon-Ultra filtration (EMD Millipore) and dialyzed against storage buffer (10 mM imidazole, pH 7.5, 300 mM NaCl, 1 mM EGTA, 1 mM NaN_3_, 50 % (vol/vol) glycerol, 1 µg/mL Leupeptin, 2 mM DTT) at 4°C.

Recombinant *S. pombe* PAK purification for phosphorylation of Myo2 (see below) was performed as previously described (Pollard et al., 2017). SNAP-Cdc12 (FH1-FH2)-His_6_ was purified using published methods (Kovar et al., 2003; Scott et al., 2011; Zimmermann et al., 2017).

### Acetylated tropomyosin

The unacetylated fission yeast tropomyosin Cdc8 was purified as previously described (Clayton et al., 2014). The acetylated Cdc8 was obtained by co-expression of Cdc8 and fission yeast N-α-acetyltransferase B (NatB) complex as described (Johnson et al., 2010) in BL21-DE3 cells. pNatB (pACYCduet-naa20-naa25) was a gift from Dan Mulvihill (Addgene plasmid # 53613). Briefly, the cultures were grown at 37 °C to reach OD_600_ of 0.8-1 and the protein expression was induced with 0.5 mM IPTG for 4-5 h at 25 °C. The cells were pelleted and sonicated in lysis buffer (100 mM NaCl, 10 mM imidazole pH 7.5, 2 mM EDTA, 1 mM DTT, 0.5 mM AEBSF, 5 µg/mL leupeptin, 0.5 mM PMSF, 5 mM benzamidine). The lysate was clarified by centrifugation, boiled for 10 min, cooled down to 25 °C, and clarified again to remove the denatured protein. The tropomyosin was precipitated by adjusting the pH of the supernatant to approximately pH 4.6. The precipitated tropomyosin was solubilized in Tpm Buffer (10 mM imidazole, pH 7.4, 50 mM NaCl, 1 mM EGTA, and 1 mM DTT) and dialyzed against the same buffer at 4 °C. The tropomyosin was then further purified on ion-exchange column with a 0.05-1 M NaCl gradient in Tpm Buffer and the proper fractions were pooled and dialyzed against Tpm buffer. To confirm the acetylation, the tropomyosin samples were dissolved in 7.5 M urea and resolved on 40 % (vol/vol) glycerol gel to compare difference in mobility (Trybus, 2000) between acetylated and unacetylated tropomyosin, based on the charge difference.

### Rhodamine labeling of actin and tropomyosin

Actin was polymerized in buffer containing 100 mM KCl, 10 mM imidazole pH 7.5, 2 mM MgCl_2_, 1 mM EGTA, 1 mM DTT at 25 °C for 2 h and dialyzed against buffer containing 50 mM PIPES (piperazine-N,N’-bis (ethanesulfonic acid)) pH 6.8, 100 mM NaCl, 0.2 mM CaCl_2_, 0.2 mM Na_2_ATP), without DTT, for 4 h to overnight. The dialyzed actin was mixed with 6 molar excess of NHS-Rhodamine (Thermo Fisher) and incubate in the dark at 23 °C for 2 h. The actin was then depolymerized and the excess dye was removed by dialysis against G-buffer for 3 d. The labeled actin was clarified by centrifugation at 350,000 g for 45 min and flash frozen in liquid nitrogen and stored at -80 °C. The percentage of labeling was calculated by the concentration of rhodamine using absorbance at 555 nm (extinction coefficient for rhodamine, 80,000 M^-1^·cm^-1^) against the protein concentration determined using Bradford protein assay (ThermoFisher).

A point mutation (D142C) was introduced in fission yeast tropomyosin Cdc8 to allow direct labeling by tetramethylrhodamine-5-maleimide (TMR-maleimide). Tpm D142C shows similar binding affinity to skeletal actin *in vitro* compared to the WT Tpm. This point mutation in Cdc8 was tolerated with modest defect on septa formation (Christensen et al., 2017). The purified Tpm (acetylated or unacetylated) was mixed with 6-fold molar excess TMR-maleimide and incubated in Tpm buffer (50 mM NaCl, 10 mM imidazole pH 7.5, 1 mM EGTA, and 1 mM DTT) at 23 °C for 1-2 h. The excess dye was removed by repeated dialysis against Tpm buffer. The labeled Tpm was stored at 4 °C up to 1 month.

### Actin polymerization visualized by TIRF microscopy

The actin polymerization assay was performed similarly as previously described (Lu et al., 2015). Actin polymer was visualized by Lifeact-GFP at final concentration of 0.82 µM. Briefly, globular actin (G-actin) and formin (when applicable) were clarified by centrifugation at 350,000 g for 30min. The concentrations of actin and other proteins in the assay were determined by Bradford protein assay (ThermoFisher) using bovine serum albumin (BSA) as a standard. G-actin is diluted to 8 µM in G-buffer as a working stock. Lifeact-GFP was diluted to 1 mg/mL in low salt buffer (10 mM imidazole pH 7.5, 50 mM KCl, 4 mM MgCl_2_, 1 mM EGTA, and 10 mM DTT) as a working stock. The glass chamber was coated with 5 µg/mL N-ethylmaleimide modified-myosin at 25°C for 2 min and blocked by 10 mg/mL BSA for 3 min, and rinsed 1-2 times with polymerization buffer (10 mM imidazole pH 7.5, 50 mM KCl, 4 mM MgCl_2_, 1 mM EGTA, 2 mM MgATP, 10 mM DTT, oxygen scavengers (3 mg/mL glucose, 0.25 mg/mL catalase, 0.13 mg/mL glucose oxidase)). Based on the final concentration of actin, 2.5-12.5 µl of 8 µM G actin is then quickly mixed with 1 µl Lifeact-GFP (1 mg/mL), G-buffer, as well as formin, profilin and tropomyosin when needed, and 19 µl 2x polymerization master mix (which gives to the final reaction in a final volume of 40 µl: 10 mM imidazole pH 7.5, 50 mM KCl, 4 mM MgCl_2_, 0.5 mM EGTA, 2 mM MgATP, 10 mM DTT, 0.25 % methylcellulose, and oxygen scavengers). Protein concentrations are indicated in each assay. The mixture was perfused into the chamber and the flowed through liquid was removed. The chamber was sealed by nail polish and imaged with ECLIPS Ti inverted microscope (Nikon) equipped with through-object type TIRF, driven by NIS Elements (Nikon). The time-lapse images were acquired by ANDOR iXon EMCCD camera through CFI Plan Apo L 100× (NA 1.45) objective at 488 nm (Lifeact-GFP), 561 nm (Rhodamine-Tpm) and 647 nm (SNAP Surface 649-labeled formin) wavelengths, at frame rates of 12-60 /min and pixel resolution at 160 nm. The focus was maintained by Ti-ND6-PFS Perfect Focus with motorized nosepiece. The polymerization was performed at 25 °C. The images were processed by ImageJ (NIH) and the actin kymographs was assembled using ImageJ with a custom plugin. The actin filaments were tracked manually. 22-60 filaments were tracked in each condition. Two independent actin preparations were used in each condition.

### Tropomyosin binding assay

3 µM of fission yeast F-actin was incubated with 0.04-4 µM of acetylated Tpm or 0.3-6 µM unacetylated Tpm at 23 °C for 10 min in binding buffer (10 mM imidazole pH 7.5, 150 mM NaCl, 2 mM MgCl_2_, 1 mM EGTA, 2 mM DTT). The mixtures were spun down at 400,000 g for 30min, the pellets were resolved on a 12 % SDS-PAGE and stained with Coomassie blue. The bands were imaged by Kodak 1D image system and the intensities of the bands were quantified by QuantityOne software (Bio-Rad). The binding constant was fit to Hill equation (Barua et al., 2013) using Graphpad Prism.

### Myo51 tail-actin pelleting assay

The assay was performed as previously described (Tang et al., 2016). Briefly, My51-Rng8/9 tail was clarified at 400,000 g for 20 min, and 1 µM of tail was mixed with 4 µM of chicken skeletal or *S. pombe* F-actin with or without 2 µM acetylated Tpm in the buffer containing 150 mM NaCl, 10 mM imidazole pH 7.5, 5 mM MgCl_2_, 1 mM EGTA, and 1 mM DTT. The mixture was incubated at 23 °C for 15 min and then pelleted at 400,000 g, the supernatant and pellet were resolved on 12 % SDS-PAGE.

### Myo2 RLC phosphorylation

Myo2 RLC was phosphorylated by incubating Myo2 with *S. pombe* PAK kinase for 1 h at 30 °C, in kinase buffer (150 mM NaCl, 10 mM imidazole pH 7.4, 5 mM MgCl_2_, 1 mM EGTA, 1 mM NaN_3_, 2 mM DTT, and 5 mM MgATP) (Pollard et al., 2017). The degree of phosphorylation was evaluated by dissolving protein samples in 7.5 M urea and resolved on 40 % glycerol gel based on charge (Trybus, 2000). For Myo2 motility assay, the phosphorylation is 100 %.

### In vitro actin motility assay

Fission yeast myosins with C-terminal biotin tag were used to allow specific attachment of the myosin tail to the assay surface as described below. The use of the actin in each step in the assay was kept species-specific to avoid mixed actin population. No phalloidin was used to keep the conditions consistent between actin species because fission yeast actin has extremely low affinity to phalloidin. Myo2-C-biotin or Myo51-Rng8/9-C-biotin was mixed with 2-fold molar excess of F-actin and 2 mM MgATP in myosin buffer (10 mM imidazole pH 7.4, 300 mM NaCl, 5 mM MgCl_2_, 1 mM EGTA, 10 mM DTT), and clarified by centrifugation at 400,000 g for 20 min to remove ATP-insensitive motor heads. The protein concentrations in the supernatant were determined by Bradford protein assay (ThermoFisher). The nitrocellulose-coated motility chamber was then coated with neutravidin (Thermo Scientific) by incubation with 0.5 mg/mL biotinylated-BSA (ThermoFisher) in Buffer A (25 mM imidazole pH 7.5, 150 mM KCl, 4 mM MgCl_2_, 1 mM EGTA, and 10 mM DTT) for 1 min followed by three washes of 1 mg/mL BSA in Buffer A, and then incubation with 10-50 µg/mL of neutravidin for 1 min followed by three washes of Buffer A. 50 µg/mL of Myo2-C-biotin or 70 µg/mL of Myo51-Rng8/9-C-biotin in Buffer A were applied to the chamber for 1 min followed by three washes of Buffer B (25 mM imidazole pH 7.5, 50 mM KCl, 4 mM MgCl_2_, 1 mM EGTA, and 10 mM DTT). The ATP-insensitive motor heads were blocked by incubation twice with 1 µM unlabeled, vortexed F-actin for 30 s each, and followed by three washes of 1-2 mM MgATP in Buffer B to release active motor heads. The MgATP was then removed by three washes with just Buffer B to allow binding to rhodamine-labeled F-actin. 60-120 nM of directly labeled actin (9-10 % labeled with rhodamine in the final concentration) was applied into the chamber twice for 30 s each. For Myo2 motility, Buffer B containing oxygen scavenger, 1mM MgATP, and 0.5 % methylcellulose was applied twice to the chamber. For Myo51-Rng8/9 motility, Motility Buffer containing 50-150 mM KCl, 25 mM imidazole pH 7.5, 4 mM MgCl_2_, 1 mM EGTA, 10 mM DTT, 70 µg/mL Cam1, 70 µg/mL Cdc4, oxygen scavenger, 1.5 mM MgATP, and 0.5-0.7 % methylcellulose was applied twice to the chamber. 2 µM acetylated Tpm was added in the Motility Buffer when actin-Tpm was tested. Actin gliding activity at 30 °C was recorded by an epifluorescences Rolera MGi Plus digital camera on a Zeiss Axiovert 10 inverted microscope equipped with a heated Plan Apo VC 60 × objective (NA 1.2), driven by Nikon NIS Element software. The movies were acquired at 2 frames/s with 0.25 µm/px resolution. Two independent actin preparations were tested in each condition. 80-300 filaments per condition of each actin were manually tracked using ImageJ (NIH) with MtrackJ plugin.

## Abbreviations

Cdc8: *S, pombe* tropomyosin
Cdc12: *S. pombe* cytokinetic formin
FH1-FH2: formin homology domains 1 and 2
For3: *S. pombe* formin
Tpm: tropomyosin

## Acknowledgements

This work is funded by R35 GM136288 to KMT, and National Institutes of Health grant R01 GM079265 to D.R.K. We thank Jenna Christensen, Dennis Zimmermann and Andrew Lombardo for technical assistance.

