## Supplemental Table 1 for "Acetylation of fission yeast tropomyosin does not promote differential association with cognate formins"

**Supporting Information**

**Table S1.** Fission yeast strains used in this study.

| Strain name | Genotype | Reference |
| --- | --- | --- |
| KV77 | h-, leu1-32, his3-D1, ura4-D18, ade6-M216 | Forsburg lab |
| KV973 | h?, act1-Y371H::kanMX6 (at endogenous locus), leu1-32, his3-D1, ura4-D18, ade6-M216 | This study |
